# A domain-general basal ganglia mechanism for inhibitory control during language comprehension

**DOI:** 10.64898/2026.11.02.763720

**Authors:** Jan R. Wessel, Yoojeong Choo, Farhad Tabasi, Cheol Soh, Nathan H. Chalkley, Héctor O. Sánchez-Meléndez, Qiang Zhang, Ergun Y. Uc, Jeremy D.W. Greenlee, Kristi Hendrickson

## Abstract

Whether controlled language processing involves domain-general or language-specific neural circuits is subject of controversial debate. We tested whether the inhibitory control of linguistic activity shares a common neural process with motor inhibition in twenty humans undergoing intra- and extracranial neural field potential recordings and deep-brain stimulation. Engaging motor and linguistic inhibition simultaneously resulted in a significant behavioral cost, suggesting a shared underlying neural process. Indeed, neural decoders trained during motor inhibition were able to decode linguistic inhibition and vice versa. Depth-electrode recordings from the subthalamic nucleus (STN), a key basal ganglia area for inhibitory control, showed increased β-band (13-30 Hz) activity during both linguistic and motor inhibition. Finally, disrupting the STN via deep-brain stimulation impaired behavioral and neural signatures of both linguistic and motor inhibition. Together, these findings show that the inhibitory control of language and movement share a domain-general mechanism implemented via the subthalamic nucleus of the basal ganglia.

## Introduction

Human language comprehension is remarkably fast and efficient, despite the often ambiguous, noisy, and widely variable nature of the incoming signal. To facilitate the real-time processing of language, the human brain constantly uses predictive processes to infer likely continuations^1–3^. At any moment, this requires selecting among competing possible words, meanings, and interpretations, while suppressing alternatives that are ultimately incorrect. This requires a substantial degree of cognitive control^1,4–7^. Whether such cognitive control processes during language comprehension are implemented by language-specific mechanisms or by domain-general executive functions is controversially debated, and remains one of the central unresolved questions in cognitive neuroscience and psycholinguistics^8–11^.

Executive functions are general-purpose control mechanisms – updating, shifting, and inhibition – that regulate the dynamics of human cognition ^12–14^. A substantial body of work in behavioral psycholinguistics suggests that controlled language processing involves domain-general executive functions that are also used to regulate non-linguistic processes. A prominent example is the transfer of control-sequence effects – improvements in executive functions after trials that already required control^15^ – across domains. Indeed, exerting executive control over non-linguistic stimuli benefits linguistic control on the next trial (and vice versa ^16,17^; but see ^18^). Moreover, individual differences in language ability and domain-general executive functions are tightly related across the lifespan, both in typical and atypical language development^19–23^. This and other work, such as observations of better domain-general executive functions in bilinguals^24,25^, has served as the basis for the argument that non-linguistic and linguistic control are governed by shared underlying mechanisms.

If that is indeed the case, linguistic and non-linguistic control should recruit a shared neural circuit. However, influential neuroscientific work contradicts this prediction. For example, an extensive series of brain imaging studies has found little overlap between the cortical networks activated during language processing and the domain-general ‘multi-demand’ network, which is associated with the executive control of non-linguistic information^8,26–29^. This absence of overlap between the neural networks for language processing and executive function has been the basis for the argument that the control of language processing is implemented in specialized, language-specific networks^30,31^.

However, existing neuroscientific work on linguistic control has two important limitations. First, the implementation of many domain-general executive functions depends on small nuclei in the human subcortex, especially those in the basal ganglia^32–36^. Many of these nuclei are too small to be imaged with the types of non-invasive methods that are typically used to investigate the role of domain-general executive functions in language processing^37–39^. Thus, it is possible that language processing and domain-general executive control may engage overlapping subcortical brain regions in the basal ganglia, whose activity may have eluded existing work. Second, the vast majority of existing work on the overlap between linguistic and non-linguistic control has been correlational (e.g., ^40,41^). Results from studies with (pseudo-)causal methods such as brain lesions have been mixed (e.g., ^42^). Notably, damage to the basal ganglia can lead to substantial impairments in language comprehension^43,44^, once again suggesting that these nuclei may be important for domain-general executive control – including that of language.

Here, we tested whether a well-characterized basal ganglia circuit for inhibitory control implements a domain-general mechanism during language comprehension. Inhibitory control is one of the primary executive functions^12,14^ and serves to suppress unwanted, outdated or inappropriate processes^32,45–47^. It is important for multiple aspects of language processing^48,49^ including the suppression of outdated semantic information^50–52^. Outside of the linguistic domain, it is well-established that the human brain exerts inhibitory control through projections from areas within the frontal cortex to the subthalamic nucleus (STN) of the basal ganglia^32,46,53–55^. The STN then exerts net-inhibitory influence on the thalamocortical drive that underpins active neural processes^56,57^. This process is best understood in the context of motor inhibition. When motor inhibition is needed, frontal cortical areas rapidly recruit the STN^58^, thereby net-inhibiting the thalamocortical drive that otherwise invigorates specific movements^59–61^. Importantly, recent work has shown that this STN-mediated inhibitory circuit is not limited to the motor domain^32^. Indeed, the STN is also involved in the inhibition of processes such as attention, decision making, and memory^62–64^. This makes the STN a prime candidate for a domain-general area that might contribute to linguistic control.

The key challenge in studying the STN is that it is a small, elongated, biconvex structure with an approximate volume of 100 mm³, which is embedded deeply in the subcortex. Non-invasive methods (such as those typically used to study language) have insufficient spatiotemporal precision to reliably measure its activity^37–39^. However, direct recordings of local field potentials from the human STN can be performed in patients with implanted deep-brain stimulation (DBS) systems, which are used to treat Parkinson’s disease (PD) and other neuropsychiatric disorders^65,66^. Recordings from DBS electrodes have greatly contributed to our understanding of STN’s inhibitory function^36,54,62^. Furthermore, DBS can be used to test the causal role of the STN. DBS produces a functional lesion of the STN and reduces its inhibitory influence^62,63,67,68^.

Here, we use intracranial recordings, multivariate analyses of simultaneously acquired scalp-EEG, and subcortical deep-brain stimulation to test whether the STN is involved in domain-general inhibitory control during language comprehension. We developed a behavioral paradigm that requires both linguistic and motor inhibition. Motor inhibition was chosen because it is the most well-documented function of the STN^32,46,53–55,58^ and was operationalized through the stop-signal paradigm^69,70^. Linguistic inhibition was operationalized via a semantic violation task, which requires the suppression of competing linguistic representations during language comprehension^50^. This is arguably the most well-documented role of inhibitory control in language processing. As linguistic input unfolds, the brain uses parallel and incremental processes to predict likely continuations, such as upcoming words in a sentence^1–3^. When those predictions are violated, the predicted word needs to be rapidly suppressed to facilitate comprehension^71–73^. Inhibitory control processes are key to this semantic suppression^7,50,51,74,75^. We hypothesized that the STN was involved in both types of inhibitory control: linguistic inhibition during the processing of semantic violations, and motor inhibition during the stopping of actions.

### Simultaneous linguistic and motor inhibition is behaviorally costly

In this combined semantic violation stop-signal task (***Fig. 1A***), highly constrained sentences were visually presented word by word. These sentences were normed for high cloze-probability – i.e., each sentence strongly primes a specific final word (e.g., *“Dad cut the bread with a …”*). Half of the sentences ended in the primed word (*“KNIFE”*), whereas the other half ended in semantic violations (e.g., *“CHAIR”*). Upon the final word (printed in all capital letters), subjects were required to quickly press one of two buttons to indicate whether the sentence was semantically sensible (*Nonviolation* trials; do not require linguistic inhibition) or not (*Violation* trials; require linguistic inhibition). On one third of trials of both types (Violation, Nonviolation), subjects were presented with a stop-signal (the final word turning red). Subjects were instructed to stop their response on those trials, and the stop-signal delay (SSD) was dynamically adjusted separately for both conditions (see ***Methods***). These *Stop* trials require motor inhibition, which was measured via stop-signal reaction time (SSRT). Twenty PD patients with implanted neurostimulators in the STN (see ***Supplementary Table 1*** for patient characteristics) performed this task.

**Figure 1.**
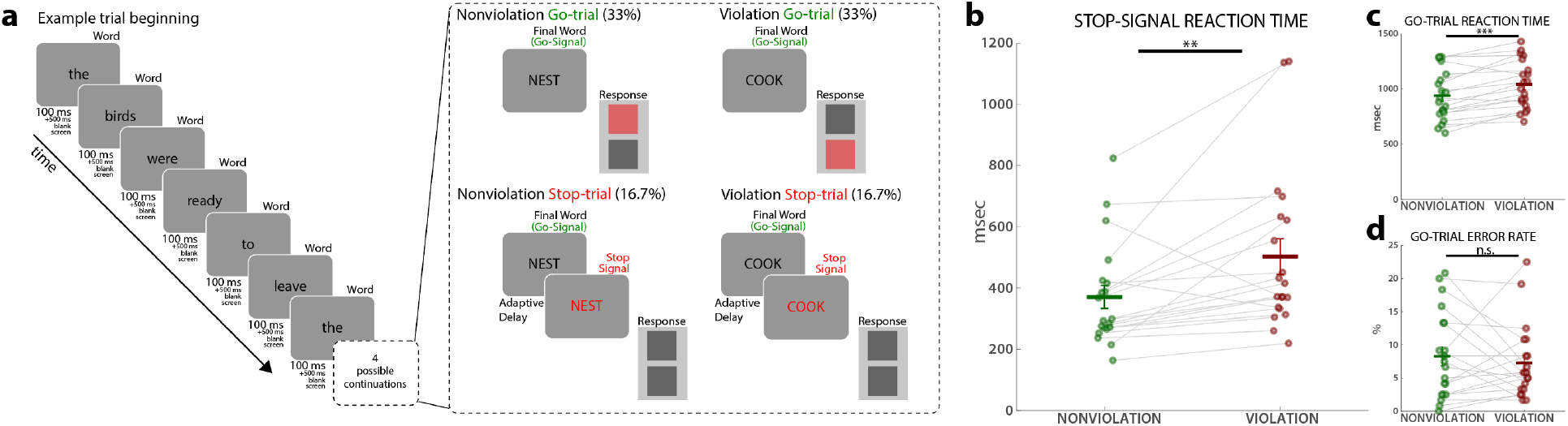
Simultaneous requirements of motor and linguistic inhibition lead to a behavioral dual-process cost, indicative of a shared underlying process. A) Diagram of the behavioral task. Participants were presented with sentences that strongly primed a specific word. The final word was presented in capital letters, which instructed participants to signal whether the sentence was sensible (i.e., the final word matched the preceding sentence context) or not (Violations). Following this ‘go’-signal, 33% of trials in both conditions featured a stop-signal, which instructed subjects to inhibit the motor response. Violation trials require linguistic inhibition, stop-signal trials require motor inhibition. Violation-Stop trials require both. B) Stop-signal reaction time (SSRT) was significantly increased on Violation trials, indicating a significant dual-task cost when motor and linguistic inhibition were required at the same time. Notably, this cost emerged despite slower responses in Violation-Go trials (c). Moreover, the impairment in motor inhibition on Violation-Stop trials was not explained by a general performance decrement on Violation trials, as go-trial error rates did not differ (d). Vertical bars denote the standard error of the mean, ** p < .01; *** p < .001.

We first hypothesized that motor inhibition would be impaired when linguistic inhibition was required at the same time (behavioral dual-process interference^76,77^). Therefore, we predicted that SSRT would be elongated on *Violation-Stop* trials (which require linguistic and motor inhibition) compared to *Nonviolation-Stop* trials (which require only motor inhibition). In line with this prediction, SSRT was significantly elongated on *Violation-Stop* (mean: 505ms, SEM: 58ms) compared *Nonviolation-Stop trials* (mean: 371ms, SEM: 38ms; t(19) = 3.13, p = .0055, d = .62; ***Fig. 1B***). This was despite the fact that subjects responded significantly slower on *Violation-Go* compared to *Nonviolation-Go* trials (1,044ms, SEM: 49ms vs. 940ms, SEM: 50ms; t(19) = 4.79, p = .00013, d = .483; ***Fig. 1C***), which should make it easier to stop the response on *Violation* trials. Importantly, the dual-process cost on *Violation-Stop* trials could not be explained by a general performance decrement, as error rates did not differ between *Violation-Go* and *Nonviolation-Go* trials (in fact, subjects were numerically more accurate on *Violation-Go trials*; error rate 7.32%, SEM: 1.26 vs. 8.31%, SEM: 1.47%; t(19) = .697, p = .494, d = .166; ***Fig. 1D;*** see Supplementary Materials for additional behavioral analyses).

Together, the behavioral results show a substantial dual-process cost on inhibitory control when both linguistic and motor inhibition were required simultaneously. This suggests that they recruit a shared neural process.

### Neural decoding analyses suggest shared neural mechanism

We then used linear discriminant classification to train multivariate neural decoders on 64-channel scalp-EEG data recorded during the task. We trained one ‘motor inhibition’ decoder on *Stop* vs. *Go* trials in the *Nonviolation* condition, which showed significant decoding from 88ms to 900ms (p < .05, cluster-corrected for multiple comparisons; ***Fig 2A***). We also trained a ‘linguistic inhibition’ decoder on *Violation* vs. *Nonviolation* trials in the *Go* condition, which showed significant decoding from 252ms to 900ms (p < .05, cluster-corrected for multiple comparisons; ***Fig 2A***).

**Figure 2.**
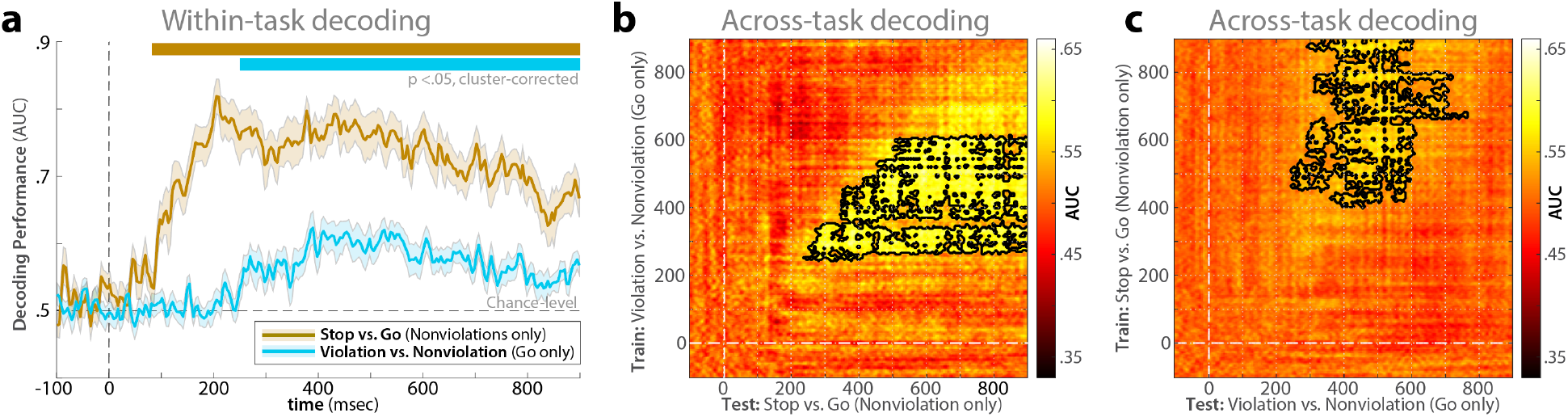
Neural decoding from scalp-EEG indicates significant processing overlap between motor and linguistic inhibition. A) Multivariate pattern analysis of whole-scalp EEG data via linear discriminant classifiers successfully decoded the condition difference for motor inhibition (Stop vs. Go trials in the Nonviolation condition) as well as linguistic inhibition (Violation-Go vs. Nonviolation-Go trials). Shaded areas represent the standard error of the decoding accuracy metric (AUC) across subjects. Colored bars at the top of the plot reflect time-periods of significant above-chance decoding (p <.05, cluster-corrected). B-C) Bidirectional cross-decoding results; the linguistic inhibition decoder was able to accurately distinguish the motor inhibition conditions (left panel) and vice-versa. Black outlines represent time periods of significant cross-decoding (p < .05, cluster-corrected for multiple comparisons).

We predicted that if motor and linguistic inhibition share a neural process, these successfully trained conditional decoders should be able to cross-decode neural activity between the condition contrasts. Indeed, the motor inhibition decoder trained to distinguish *Nonviolation-Stop* from *Nonviolation-Go* trials was able to successfully decode the linguistic inhibition condition contrast between *Violation-Go* and *Nonviolation-Go* trials (p < .05; cluster-corrected for multiple comparisons) and vice versa (***Fig. 2B***). Importantly, the motor inhibition decoder did not contain instances of linguistic inhibition (i.e., no *Violation* trials were included in training the motor inhibition decoder) and the linguistic inhibition decoder did not contain instances of motor inhibition. Moreover, contrasts with overlapping conditions used different trials from each condition. Finally, to ensure that this successful cross-decoding was not attributable to motor differences in the linguistic inhibition contrast, we also repeated this analysis after removing the neural independent components underlying the motor response (***Fig. S1A***). Cross-decoding remained significant; indeed, cross-decoding accuracy numerically increased (***Fig. S1B***).

### STN activity is increased during both motor and linguistic inhibition

We then tested whether local field potential activity in the STN was increased during both motor inhibition (which is well established^54,55,60,78–80^) and linguistic inhibition. Local field potentials from STN were recorded from bilaterally implanted depth electrode probes (***Fig. 3A***) using a novel telemetry-based technique that allows post-surgical recordings from the STN^81^. In line with prior work, we found significant increases in STN β-band (13-30 Hz) activity on Stop vs. Go trials (***Fig. 3B, bottom row***). 18 out of 20 participants showed such increased β-band activity during the significant time-frequency cluster in right STN, and 16 out of 20 participants showed increased β in left STN.

**Figure 3.**
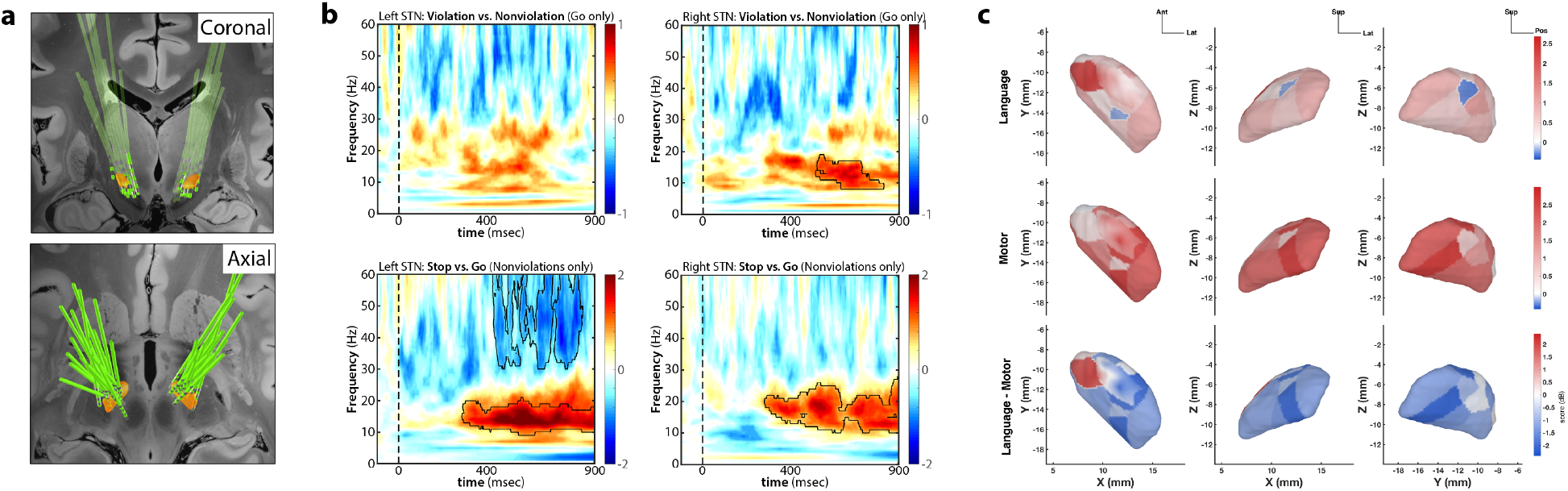
Subthalamic local field potentials during motor and linguistic inhibition. A) Implantation trajectories (green) and electrode contact locations relative to STN (orange); coronal (top) and axial (bottom) views. B) Event-related spectral perturbation during linguistic inhibition (top row) and motor inhibition (bottom row), separately for left STN and right STN. In line with prior work, motor inhibition was marked by significant bilateral increases in β band (13-30 Hz) activity on Nonviolation-Stop vs. Nonviolation-Go trials. Similar β band increases were observed for linguistic inhibition (Violation-Go vs. Nonviolation-Go trials). After correction for multiple comparisons, these remained significant in right STN only. However, comparisons between left and right STN were not significant (**Fig. S2**). Black outlines demarcate significant condition differences at p < .05, cluster-corrected for multiple comparisons. C) Topographical distribution of β band activity in STN, separately for language contrast (top row), motor contrast (middle row), and difference (bottom row).

Crucially, β-band activity was also increased during linguistic inhibition (***Fig. 3B, top row***). After correction for multiple comparisons (p < .05, cluster-corrected), this group-level increase remained significant in the right STN. On the individual level, 17 out of 20 participants showed a β band increase during the significant time-frequency cluster in right STN. The left STN showed the same numerical activity increase as right STN, but this did not survive correction for multiple comparisons. We had no a priori hypothesis regarding lateralization. Direct comparisons between right and left STN did not yield any significant differences in either contrast (***Fig. S2***).

***Figure 3C*** shows the spatial distribution of the significant β-band activity increase in right STN for both motor and linguistic inhibition, as well as their spatial difference topography. The activity increase during motor inhibition showed a linear spatial organization (*R*^2^ =.46; F(3,13) = 3.72, *p*perm = .034), with a cross-validation predictability LOO-*R*^2^ = .15. No single site drove this model (refitting the data after exclusion of a site that showed a Cook’s distance above the critical threshold of D = .25 left the model fit virtually unchanged, *R*^2^= .48). Across all 17 refits, *R*^2^ ranged from .39 to .58. This spatial organization trend was mainly observed along the mediolateral axis, with larger responses at more lateral sites (βX = .35; βY = -.02; βZ = -0.15). Spearman correlation coefficients showed the same directional organization pattern (⍴ = .66, -.52 and -.43 for X, Y and Z, respectively). This is in line with previous work showing stronger involvement of dorsolateral STN sites for motor inhibition (e.g., ^82^). In contrast, language effects were more diffusely distributed. The linguistic inhibition contrast showed no evidence of a linear spatial organization (*R*^2^ = .16, adjusted-*R*^2^ = -.03, F(3,13) = .83, *p*perm = .49; LOO-*R*^2^ = -0.27).

### Functional lesions of STN impair motor and linguistic inhibition

The depth-electrode recordings showed that the STN is active during both motor and linguistic inhibition. To provide converging causal evidence for the proposition that the STN contributes to both types of inhibition, we had all patients perform the task twice, once with DBS *On* and once with DBS *Off* (with counterbalanced session order). All of the above results were from the *Off* session, when STN function is not altered by DBS. We compared behavior and scalp EEG activity between the *On* DBS session (when STN activity is disrupted) and the *Off* session. STN DBS is known to reduce motor inhibition^67,83,84^. Here, we tested whether the same was true for linguistic inhibition. To ensure that the DBS electrodes were within language-related regions of the STN, these analyses were restricted to the 17 participants in which the right STN contact showed increased STN β band activity in the language contrast in ***Fig. 3B***. In line with our hypotheses, DBS impaired both motor and linguistic inhibition. Behaviorally, DBS elongated SSRT (mean effect of DBS: F(1,16) = 6.59, p = .021, η^2^ = .29; ***Fig. 4A***), as has been previously found (e.g., ^83,84^). Crucially, there was a significant interaction between DBS condition and linguistic inhibition (F(1,16) = 10.43, p = .0052, η^2^ = 0.39) – in other words, STN DBS significantly increased the behavioral cost that occurred when both motor inhibition and linguistic inhibition were required (***Fig. 4A***). Importantly, these effects were specific to inhibition, as DBS did not affect basic *Go* reaction time (main effect of DBS: F(1,16) = .0261, p = .874, η^2^ = .0016) and did not alter the reaction time difference between *Violation-Go* and *Nonviolation-Go* trials (DBS x Violation interaction: F(1,16) = .119, p = .734, η^2^ = .0074). Similarly, DBS also did not affect RT on failed Stop trials (main effect of DBS: F(1,16) = .497, p = .491, η^2^ = .0301; DBS x Violation interaction: F(1,16) = 1.45, p = .246, η^2^ = .083) or error rate (main effect of DBS: F(1,16) = .209, p = .653, η^2^ = .0129; DBS x Violation interaction: F(1,16) = .454, p = .51, η^2^ = .0276).

**Figure 4.**
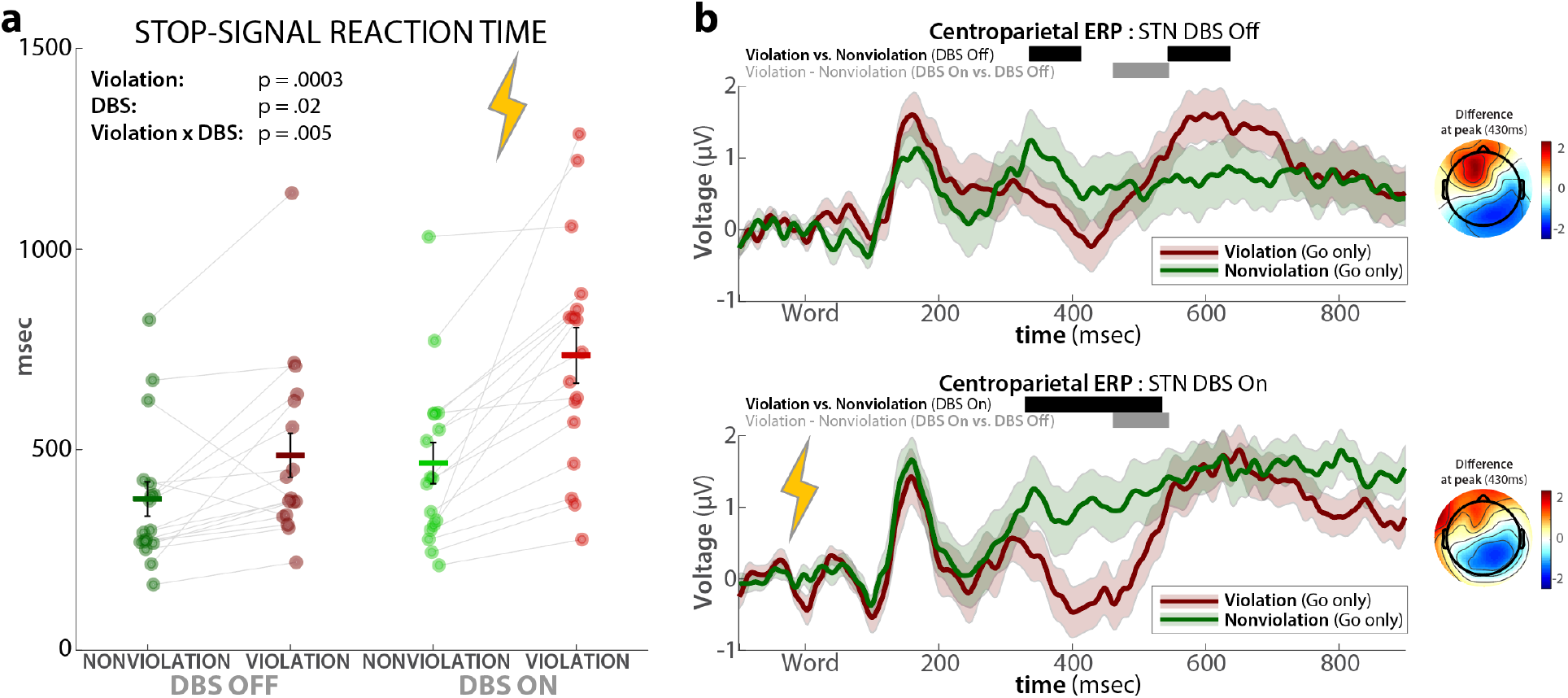
Effects of STN DBS on behavioral and electrophysiological indices of motor and linguistic inhibition. A) STN DBS increased stop-signal reaction time and showed a significant interaction with linguistic inhibition (greater SSRT increase on Violation trials). Vertical bars indicate the standard error of the mean. B) STN DBS increased the N400 event-related potential. Off DBS event-related potential on top, On DBS event-related potential at the bottom. Shaded areas reflect the standard error of the mean. Black horizontal bars denote significant condition difference between Violation and Nonviolation trials within each DBS condition (p < .05, cluster corrected for multiple comparisons). Gray bars denote significant differences in the Violation-Nonviolation difference wave between the DBS conditions (p < .05 cluster-corrected for multiple comparisons). Voltage topographies show the scalp distribution of the difference wave, separately for On and Off DBS, at the peak latency of the respective difference wave (430ms in both On and Off DBS conditions).

To further assess whether DBS affected language-related processes, we tested whether DBS significantly altered the amplitude of the centro-parietal N400 event-related potential (***Fig. 4B***). The N400 is the primary event-related potential investigated in semantic violation tasks. Its amplitude indexes the degree of difficulty associated with integrating the word into its semantic context (see ***Discussion***). A significant N400 ERP was observed both *On* and *Off* DBS (p < .05, cluster-corrected; black bar in ***Fig. 4B***). Crucially, the amplitude of this N400 was significantly increased *On* DBS (p < .05, cluster-corrected; gray bar in ***Fig. 4B***), suggesting that disrupting linguistic inhibition implemented by the STN increases the difficulty of integrating the violating stimulus into the semantic context.

Together, these findings demonstrate that DBS causally affects both motor and linguistic inhibition.

## Discussion

We identify a domain-general neural mechanism for inhibitory control during language comprehension, implemented via the subthalamic nucleus of the basal ganglia. This crucially extends recent work on the domain-general inhibitory role of the STN^35,36,62–64^ to arguably the most uniquely human ability – language. These findings explain why previous neuroscientific work has largely failed to identify overlap between brain networks for executive function and language processing, despite this being suggested by much behavioral work: basal ganglia regions like the STN are too small to be reliably imaged using the non-invasive methods that have been used in existing neuroscientific work on controlled language comprehension^30,38,39^. Here, we used a new telemetry-based LFP recording technique that enables intracranial recordings from chronically implanted electrodes^81,85^. This technique overcomes the typical limitations of human basal ganglia recordings, which typically have to be performed during awake neurosurgery. Instead, the current recordings were performed months or years after successful surgery, when DBS treatment has been optimized, with normally awake and alert participants on their usual dopaminergic treatment. This allowed us to collect extensive LFP and behavioral data in each patient (50+ minutes per session), collect an extraordinarily large sample (N = 20) compared to existing intracranial recording studies from the human basal ganglia, measure simultaneous high-density scalp EEG to quantify the N400 ERP, and have the same participants perform multiple sessions to test the causal role of the STN *On* vs. *Off* stimulation.

Evidence for the overlap between linguistic and motor inhibition converged across all four levels of analysis. First, carrying out linguistic and motor inhibition at the same time was behaviorally costly, resulting in elongated SSRT during semantic violations. Such behavioral dual-process costs are a well-established indicator of a shared underlying process^77,86^. Indeed, motor inhibition can be paired with other tasks without incurring a cost on SSRT when the underlying processes do not overlap (e.g., ^87^). Second, multivariate neural classifiers trained on motor inhibition could be used to decode linguistic inhibition (and vice versa), further suggesting a shared process that is active in both conditions. Direct LFP recordings then showed that STN β-band activity was significantly increased during both motor and linguistic inhibition. This is in line with past work on the role of STN β-band activity in motor inhibition^54,55,60,78,80,88^ and in inhibitory control more broadly^32,89^. Finally, we causally disrupted the STN via DBS, which reduces its inhibitory function^67,83,84^. This led to an elongation of SSRT that was further increased in the linguistic inhibition condition, as well as an increase in the N400 ERP.

The involvement of the STN in linguistic inhibition is consistent with its proposed role as a truncator of thalamocortical drive^32^. This mechanism is well-established during motor inhibition, where frontal cortical areas rapidly recruit the STN^58^, which then net-inhibits the thalamocortical drive that otherwise invigorates movements^59–61^. Just like movement, language processing involves the thalamus^90,91^. Notably, past recording work has shown that language-related activity in the thalamus changes significantly during semantic violations^92^. Our findings provide a mechanistic explanation for this activity change, suggesting that the STN inhibits language-related thalamocortical drive during semantic violations. Indeed, when STN was disrupted via DBS, participants exhibited both a larger behavioral dual-task cost (***Fig. 4A***) and increased the N400 ERP amplitude to semantic violations (***Fig. 4B***). The latter finding is especially interesting. Influential accounts of the functional significance of the N400 suggest that it indexes the difficulty of semantic access or the difficulty of integrating word meaning into its context^93,94^. Past work has hypothesized that inhibition of the initially predicted word aids this process when semantic violations occur^50–52^ (see also ^95^). Under this account, reducing the ability to exert linguistic inhibition (here, by disrupting the STN) increases the difficulty of semantic access or integration, leading to a larger N400 semantic effect. The larger N400 semantic effect observed with DBS *On* is consistent with this account. This account may also help explain some surprising clinical findings regarding the effects of STN DBS on language comprehension in Parkinson’s Disease. While language comprehension is typically unimpaired in PD, patients treated with STN DBS can show language comprehension deficits that are absent when the DBS is inactive ^96,97^. The current work suggests a possible mechanism: DBS disrupts the inhibition needed to facilitate language comprehension, especially during situations that require control.

What other aspects of language comprehension are aided by the STN, or by domain-general neural processes in general, should be the subject of future study. The current work provides a first description of a specific, domain-general neural circuit for controlled language processing during semantic violations. Other types of control-demanding language comprehension processes might recruit this same circuit. For example, syntactic violations activate a fronto-subcortical network that overlaps substantially with the motor inhibition network^98^, suggesting that the same domain-general inhibitory control network may be involved. The processing of negations is another likely candidate. Negations feature substantial activation of the (negated) concept or statement, thereby creating competition that may require inhibition to properly resolve. Indeed, negations recruit several brain regions that are also active during motor inhibition^99^. However, other forms of linguistic competition are less likely to involve this inhibitory circuit. Competition during word recognition, for example, may be resolved through local inhibitory interactions among lexical representations, as proposed by connectionist models of word recognition^2^. Future studies are necessary to directly test these hypotheses.

Finally, we used precise 3D reconstructions of the STN recording locations in each patient to explore a potential subregional organization or lateralization of language- and motor-related activity within STN. At the cortical level, language is mostly implemented in left-lateralized regions, and the motor system is organized contralaterally to the respective effector(s). However, recent work on the human STN found that inhibitory control – at least of the motor system – is implemented through synchronous, bilateral STN activity, regardless of the side of the motor effector^80^. Similarly, we here found no evidence for a systematic lateralization of linguistic inhibition activity at the level of the STN. With regards to the STN’s subregional organization, influential work in non-human animals has suggested that there is a spatial gradient in both connectivity and function of the STN^100^. However, it is debated whether there is a similarly strong spatial segregation of subregions within the human STN^101,102^. In line with previous work^82^, we found that motor activity was stronger in (dorso)lateral regions of the STN, activity related to linguistic inhibition was more diffusely distributed. Strong inferences regarding the subregional organization of STN with respect to language will necessitate systematic investigation with newly-available high-density recording arrays^103^.

Taken together, the current work shows that the STN implements a domain-general mechanism for both motor and linguistic inhibition. This extends the effective range of this well-known subcortical mechanism for inhibitory control to what is arguably the most human ability, offers a unifying explanation that bridges the gap between previously incommensurable findings in behavioral psycholinguistics and cognitive neuroscience, and contributes to our understanding of the (side)effects of STN DBS as a treatment of neuropsychiatric disorders.

## Methods

### Data and code availability

All deidentified raw data and analysis code will be made publicly available on Zenodo upon acceptance of the final manuscript.

### Participants and procedure

Participants were recruited from JDWG’s neurosurgical practice. All experiments were clinically supervised by an attending neurologist with subspecialty training in Parkinson’s disease (EYU or QZ). Participants signed a consent form approved by the University of Iowa IRB board (IRB ID: 202002751). Enrollment in the study was contingent upon scoring greater than 19 on the Montreal Cognitive Assessment (MoCA) and screening by an attending neurologist with subspecialty training in Parkinson’s disease (EYU). We recruited twenty-four Parkinson’s Disease patients who had been previously implanted with wireless sensing capable *Percept* PC DBS stimulators (*Medtronic plc., Minneapolis, MN, USA*) targeting the bilateral STN. Participants completed two experimental sessions with the DBS either turned *On* or *Off*, in randomized order. STN DBS was applied through the typical clinical contacts in the *On* session, at each participant’s typical clinical settings. This ensured a manipulation check of DBS efficacy (via the clinical benefit on UPDRS scores). Four participants elected not to return for the second visit, leaving a final sample of N = 20. The average interval between sessions was 8.1 days (Minimum interval: 7 days). Three patients had stopped taking dopaminergic Parkinson’s Disease medications due to the effectiveness of the DBS treatment. The remainder of the patients performed the task on their routine medications to allow for ideal cognitive and motor performance. (Mean Age: 65.7; Age Range: 49-80; 4 female; all right-handed; for clinical characteristics see **Supplementary Table 1**). All experiments were conducted in accordance with the Declaration of Helsinki and approved by the University of Iowa Institutional Review Board (IRB # 202002751). Stimuli were presented using Psychtoolbox 3^104^ using MATLAB 2021a (TheMathWorks, Natick, MA) on Ubuntu Linux computers connected to a BenQ XL2420B monitor featuring a 120 Hz refresh rate and a resolution of 1920x1080 pixels. Responses were made using a standard QWERTY keyboard.

### Experimental task

The task is a combination of a stop-signal task and a semantic Violation task. Participants were presented with high-constraint sentences, one word at a time. 45 Sentence stimuli were taken from a published set of sentences with established cloze probability and completion norms^105^. All sentences were nine words in length, with an average cloze probability of .90 (range .81 -.98). To make the Violation sentences, the 45 sentence frames (the first eight words) were paired with final words of other sentences. To avoid any phonological, semantic, or associative overlap between the Violation words and the high-cloze targets 12 of the final words could not be used as violating words and were replaced with other sentence final words from the completion norms database (see ^52^). Words were presented centrally in black font on a gray background. Each word was presented for 100ms, followed by a 500ms blank screen. The final word of the sentence was presented in all caps.

There were two experimental sentence conditions. In the Nonviolation sentence condition, the final word was the predicted, high-cloze word primed by the sentence context. In the Violation sentence condition, the final word was a semantic Violation. Each sentence was presented eight times, four times with the violated and four times with the non-violated ending, resulting in 360 trials per session. Participants were instructed to indicate whether the sentence was sensible (Nonviolation) or nonsensical (Violation) as fast as possible after presentation of the final word by pressing either the CAPS LOCK or SHIFT keys on the keyboard, respectively (the mapping of button to response was randomized across subjects). Responses were made with the patients’ preferred hand (typically the less-affected side of the body; 10 chose left and 10 chose right). A vertical key arrangement was chosen to avoid potential interactions with reading direction. The final word remained on the screen until a response was made. A response deadline was set at 2,000ms.

The stop-signal component was implemented by introducing the requirement that on one third of trials in both conditions, a stop-signal was presented. On those trials, the final word changed from black to red after an adaptive delay (the stop-signal delay, SSD). Participants were instructed that they had to attempt to abort the response on those trials. The SSD was initially set to 250ms and then adjusted after each stop trial; after successful stop trials, it was increased by 50ms, whereas after failed stop trials, it was decreased by 50ms. The SSD was tracked separately for the two semantic conditions (Violation and Nonviolation). This ensures that stopping is consistently challenging and results in a probability of successful stopping of ∼.5 in both conditions, which facilitates the computation of SSRT.

Feedback message appeared 500ms after the response deadline and remained on the screen for 500ms. Correct responses on Go trials and correctly withheld responses on Stop trials were followed by “Correct” and “Correct Stop”, respectively. Misses (no response within deadline window on GO trials) were followed by a “Faster!” message, whereas Errors (wrong response on GO trials) were followed by “Incorrect”. Finally, failed Stop trials (Stop trials with a response) were followed by “Please stop”. After this feedback and a 500ms blank screen inter-trial interval period, the next trial began.

The 360 total trials were divided into eight blocks of 45 trials. Trials were presented in pseudo-random order, with two constraints: the maximum number of Stop trials in a row was kept to 4, and the maximum proportion of Go trials per block was .4.

### Behavioral analysis

The dependent variable of primary interest was stop-signal reaction time (SSRT; ^69^). SSRT was calculated using the integration method with replacement of Go-trial misses^70^. Secondary parameters of interest were go-trial RT, go-trial error rate, SSD, and failed-stop trial RT. Statistical comparisons were made using *paired-samples t-tests* comparing *Violation* and *Nonviolation* trials. For the comparison of DBS conditions, a 2x2 repeated-measures ANOVA with the factors DBS (*On*, *Off*) and Violation (*Violation*, *Nonviolation*) was performed.

### LFP and EEG recording

These procedures were implemented as described in Soh et al.^80^; the description is adapted from therein. The *Percept* PC Neurostimulator’s communicator device (Medtronic model 8880T2) was used to wirelessly stream STN LFP data to the provided programmer tablet (Model CT900, *Samsung Galaxy Tab S5e*) via Bluetooth at 250 Hz (the maximum available frequency). The two directional sensing DBS leads are equipped with four rows of depth electrodes each, spaced 1.5 mm apart. LFP recordings were made from bipolar contact montages above and below the clinical stimulation electrode(s) on each side, thus centering the recorded voltage on the clinically efficacious contact site. The data were online filtered using a lowpass filter of 100 Hz and a highpass filter at 1 Hz. To ensure a stable wireless connection, both the communicator and the operating tablet were consistently kept within a 50 cm range. LFP recordings were performed with the DBS stimulator in the off state to enable the wireless sensing feature. An attending neurologist verified the DBS parameters before and after the experiment to ensure that participants left with their original DBS configurations and their DBS enabled.

EEG was acquired using a 64-channel EEG cap, amplified via Brain Vision MRplus amplifiers (*Brain Products, Garching, Germany*). The sampling rate was set to 5,000 Hz to enable a precise alignment between EEG and LFP data and to ensure sufficient sampling rate for EMG data acquisition. The EEG signal was online filtered at a low cutoff .01 Hz and a high cutoff of 1000 Hz. The reference electrode was Pz, and the ground was Fz.

Before each block of the task was started, the DBS was activated for ∼12 seconds and then turned off again. The next block did not start until 12 seconds after the DBS was turned off. This procedure introduced simultaneous artifacts in both the EEG and LFP traces, which were used to precisely align the EEG and LFP datasets via the PEA toolbox^81^. After successful alignment, the LFP data were analyzed using the digital event information from the EEG system.

### EEG preprocessing

Data were preprocessed using standard procedures via custom routines in MATLAB and the EEGLAB toolbox^106^. The data were filtered using symmetric two-way least-squares finite impulse response filters (high-pass cutoff: .3 Hz, low-pass cutoff: 50 Hz). Non-stereotyped artifacts were automatically removed using segment statistics applied to each second-long segment of data (joint probability and kurtosis, both cutoffs = 5*SD, ^107^). The resulting data were decomposed via a temporal infomax independent component analysis with a 40-dimensional input matrix enforced by principal component analysis. The resulting component matrix was visually screened for components representing eye-movement and electrode artifacts, which were removed from the data.

### Multivariate EEG decoding

Multivariate pattern analysis of the whole-scalp EEG activity was performed via the MVPA-Light toolbox^108^. The preprocessed data were downsampled to 250 Hz and segmented around - 100 to 900ms with respect to the stimuli of interest. Each epoch was baseline corrected with the 100ms pre-stimulus window. We trained and tested two multivariate classifiers for pair-wise comparisons. A Nonviolation-Stop vs. Nonviolation-Go decoder was trained to distinguish EEG activity related to motor inhibition, in the absence of linguistic inhibition demands. Conversely, a Violation-Go vs. Nonviolation-Go decoder was trained to distinguish EEG activity related to linguistic inhibition, in the absence of motor inhibition demands. Since go-trials outnumber stop trials by a factor of 3:1, we paired each successful stop-trial with the closest go-trial at the same SSD (sampling without replacement). In case the EEG preprocessing resulted in minor trial number differences between the Violation and Nonviolation conditions (which were otherwise equiprobable), we included only the N first instances of the more common condition (with N being the number of trials in the less common condition). A linear discriminant classifier was then trained and tested at each time point for each subject separately, using a four-fold cross-validation procedure. Trials from each subject were randomly separated to four sets, with one set held out as a test set and the remainder of the data used as training sets. This procedure was iterated four times until each set served as a testing dataset. The procedure was repeated five times and the resulting decoding performance was averaged across each repetition. Each subject’s decoding performance was then averaged together to be used in group-level analysis. We used the area under the curve (AUC) metric to evaluate the quality of decoding performance for both decoders.

To assess statistical inference, each participant’s AUC per time point was submitted to paired-samples Wilcoxon signed-rank test against chance-level performance (.5). The resulting *p*-values were corrected for multiple comparisons using cluster-based permutation tests (1000 iterations, *p* < .05, ‘maxsum’ cluster correction^109^).

We then performed a cross-decoding analysis^110^, wherein a decoder trained on one condition difference (e.g., the motor inhibition decoder trained on the difference between Nonviolation-Stop and Nonviolation-Go trials) was tested for its ability to decode another condition contrast (e.g., Violation-Go vs. Nonviolation-Go trials). Since both decoders included the Nonviolation-Go condition, we ensured that both training sets included a different set of Nonviolation-Go trials. The segmented EEG data were first z-scored. A time-by-time temporal generalization matrix was then obtained for each subject. As this process is stochastic (due to random assignment of trials for each dataset), we repeated this process 10 times and averaged the results to derive a single, reliable generalization matrix for each subject. Each participant’s decoding performance in this time-by-time matrix was then submitted to Wilcoxon sign-rank test against chance level performance. Multiple comparisons were again corrected with cluster-based permutation test with the critical cluster *p*-value of .05 (1000 iterations, p < .05; weighted cluster mass correction; weight = 1^109^).

### STN electrode reconstruction

DBS electrode locations were reconstructed using LeadDBS (v3.2)^111^ and Lead Group^112^, as explained elsewhere^80^. We used pre-surgery and post-surgery computed tomography (CT) and magnetic resonance (MR) images from 17 out of 20 recruited subjects (the requisite postop imaging was not available in the remaining three). Postop CTs were aligned to preop MR images for each patient using the Advanced Normalization Tools (ANTs)^113^. Normalization and brain shift were visually inspected and corrected if needed, and then normalized to MNI space (ICBM 2009b NLIN asymmetric)^114^ using ANTs symmetric diffeomorphic registration (ANTs SyN)^113^ or, alternatively, the Unified Segmentation approach^115,116^. The refined TRAC/CORE method was used to automatically reconstruct DBS contacts and was further adjusted manually. This cohort used two types of DBS electrodes: directional Medtronic B330150 and non-directional Medtronic 3380 (Medtronic, MN, USA); corresponding lead models were adjusted accordingly. Lead models for each subject, their active contact, and MNI coordinates are summarized in ***Supplementary Table 2***. Further, STN segmentation was defined by the DISTAL atlas^117^. Additionally, patients’ native anterior commissure-posterior commissure (ACPC) coordinates from clinical notes were used to cross-validate the deepest contact position. Cohorts’ DBS leads are shown relative to the subthalamic nucleus (STN) location on a 7T ex-vivo 100 μm brain atlas in ***Figure 3***^118^.

### LFP analysis

Full-spectrum event-related spectral perturbation (ERSP) plots were generated by transforming the LFP time series into the time-frequency domain via the filter-Hilbert method, with individual frequencies ranging from 1 to 60 Hz in linear steps of 1 Hz and a range of +/-.5 Hz around each center frequency. The resulting complex time series was converted into power units through squaring its real part. Event-related time-frequency activity was then extracted from - 100ms to 900ms around each event of interest and converted into change from baseline (decibel) using the 100ms prior to the event as a baseline. ERSPs were tested for significance via nonparametric permutation testing (5,000 iterations), cluster-corrected for multiple comparisons (voxel-wise p = .05, cluster-wise p = .05) across the entire time-frequency spectrum.

### Spatial mapping of LFP activity in STN

We extracted the mean LFP amplitude in the significant cluster of right STN activity for the motor and linguistic (*v*1) and motor (*v*2) contrasts in ***Fig. 3B*** for each subject. We then modeled the MNI coordinates of the recording site using ordinary least squares (MATLAB *fitlm*):

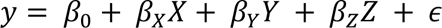

where *β* is the coefficient and *ε* is the error, to test whether the response varies linearly with the location relative to the STN. Each covariate tests the unique contribution of each axis while controlling for the other.

Since both measures, language and motor scores, are condition differences in dB, to test whether spatial differences between motor and language vary across the STN, we directly tested paired differences:

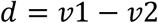

where v1 is the language, and v2 is the motor observation, and then fit

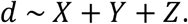

We then used a permutation test to validate the overall spatial model by randomly shuffling the response values among locations for 10,000 iterations, while anatomical coordinates are fixed, and refitting the model each time:

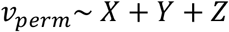

and comparing the observed F-statistic against the permutation distribution. The p-value was defined as

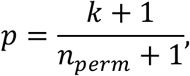

where *k* is the number of permutations whose F-statistic is at or above the observed F. Axis contributions within each multivariate model were tested using Freedman-Lane residual permutation tests (10,000 iterations). Spearman correlations between each response and each coordinate were calculated for exploratory purposes.

To test the robustness of findings, we used a leave-one-out (LOO) approach to evaluate 1) stability of the model and 2) predictive accuracy by removing each site in turn and refitting the model. Cross-validation *R*^2^*is* was calculated as:

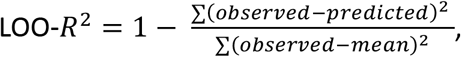

where LOO-*R*^2^ ≤ 0 indicates no out-of-sample prediction, and root-mean-squared prediction error.

### ERP analysis

The preprocessed EEG data for the DBS *On* and *Off* sessions were epoched from -100ms to 900ms relative to final word on Violation-Go and Nonviolation-Go trials and then averaged for each condition. The N400 is a negative polarity ERP that is increased on Violation compared to Nonviolation trials. The N400 was quantified at centro-parietal electrode CP2 based on the peak of the voltage difference topography. Condition differences within each session (Violation-Go vs. Nonviolation-Go) were tested using nonparametric permutation testing (5,000 iterations), cluster-corrected for multiple comparisons (voxel-wise p = .05, cluster-wise p = .05) at each time point following word onset. To test whether DBS modulated the N400, we tested the Violation-Go minus Nonviolation-Go difference wave between the DBS conditions using the same statistical approach.

## Notes

### Competing Interest Statement

The authors have declared no competing interest.

### Funding

- National Institutes of Health: R01 NS117753

