## Supplementary Materials for "A domain-general basal ganglia mechanism for inhibitory control during language comprehension"

**Additional behavioral results**

Due to the adaptive thresholding procedure for the stop-signal delay (which independently updated the stop-signal delay to achieve an approximate p(stop|signal) of .5 on *Stop* trials of both conditions), there were no differences in stopping success rate between *Violation* and *Nonviolation* trials ($t(19) = .04$, $p = .98$, $d = .007$). Nor were there differences in mean SSD ($t(19) = .348$, $p = .732$, $d = .077$). Reaction times on failed stop-trials were faster than on go-trials for both *Violation* ($t(19) = 5.17$, $p < .0001$, $d = .521$) and *Nonviolation* ($t(19) = 6.09$, $p < .0001$, $d = .697$) trials, which is a requirement for SSRT computations via the horse-race model (see Methods; Logan et al., 1984).

**MVPA Decoding after removal of independent EEG components reflecting motor activity**

Motor independent components were identified as described in (Soh et al., 2026), the description is adapted from therein. EEG signal components reflecting motor activity were identified using an automated algorithm. Each independent component from each subjects' ICA solution were backprojected into channel space. The resulting activity at motor electrode sites (C3 and C4) was then converted into β burst probability time series using established methods (Soh et al., 2026). The resulting time series were then epoched with respect to the stimulus onset on correctly-responded Go trials and then baseline-corrected with respect to the preceding 500ms period. The 200ms pre-response period for each trial was then saved and averaged across trials. The components with the largest reduction of β bursting in this time period, relative to pre-stimulus baseline, were then selected as motor components, separately for left (C3) and right (C4) motor cortex.

***Fig. S1A*** shows the features of the thusly selected components. The left panel shows beta burst probability timelines for the electrode contralateral to the respective subjects' response-hand, time-locked to the Go-signal on Nonviolation trials. Orange lines are individual subjects, black line is the subject average. Clear movement-related β suppression is evident in all subjects. The inlet shows the component weight matrix for the selected independent component. For subjects who responded with their left hand, we left/right flipped the weight matrix before averaging to rectify the average topography between subjects. A clear hotspot over left motor cortex electrode C3 is visible. The right panel shows the trial-wise beta burst probability timelines, sorted by RT. The β suppression is clearly aligned to movement.

***Fig. S1B*** shows the same MVPA decoding analysis as in the main manuscript (Fig. 2), but after removal of the independent signal components described above. Cross-decoding remained significant, and actually increased numerically.

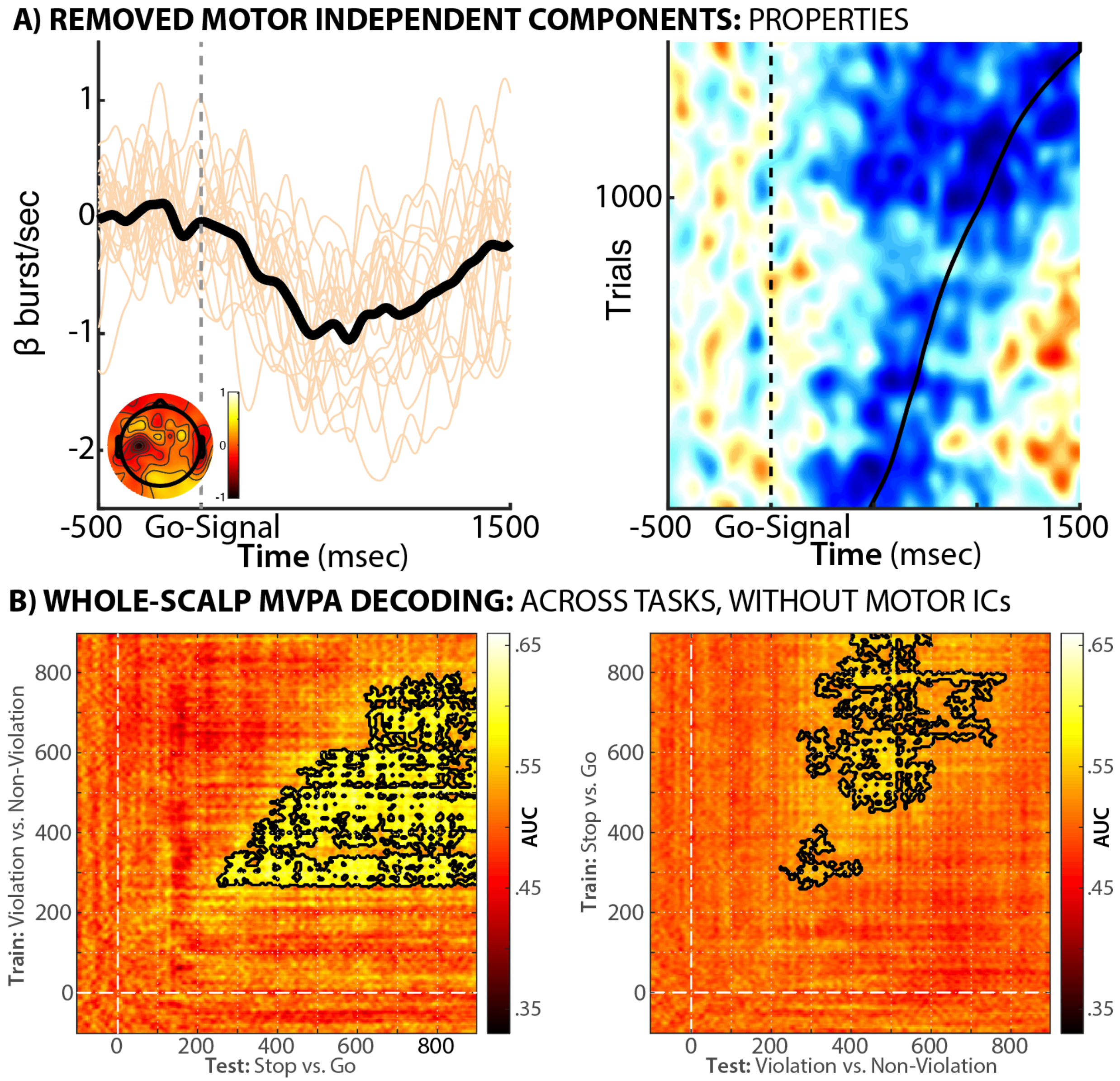


***Figure S1.*** *A) Properties of the selected independent components that reflect motor processes. B) Bidirectional cross-decoding results after removal of the independent components reflecting movement; the linguistic inhibition decoder was able to accurately distinguish the motor inhibition conditions (left panel) and vice-versa. Black outlines represent time periods of significant cross-decoding (p < .05, cluster-corrected for multiple comparisons).*

**No lateralization of STN effects for motor and linguistic inhibition**

The below comparisons show the difference between left and right STN for the two main contrasts of interest (Violation-Go vs. Nonviolation-Go and Nonviolation-Stop vs. Nonviolation-Go). There were no significant differences for either comparison (p < .05, cluster-corrected).

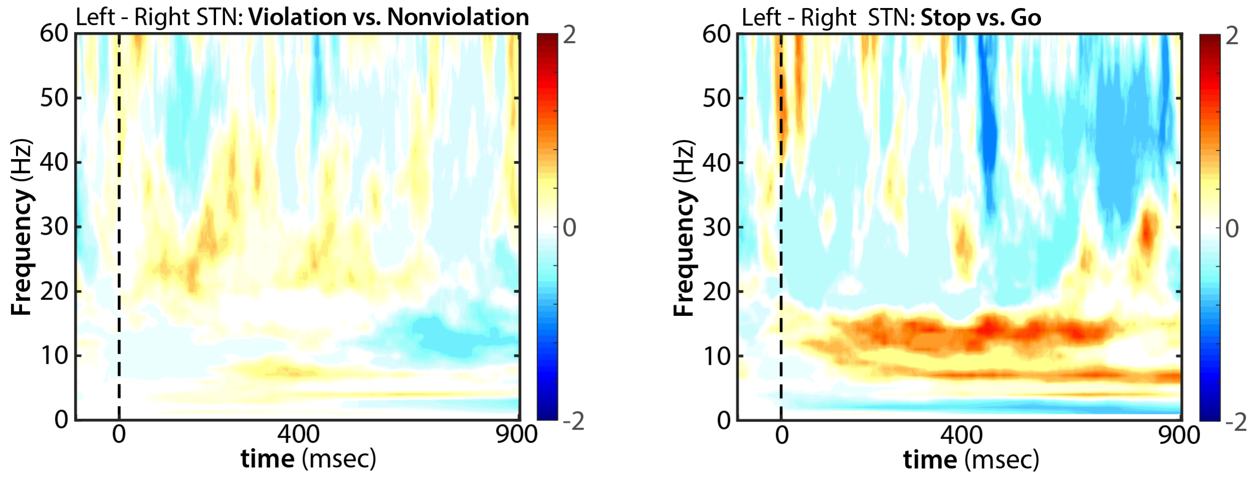


***Figure S2.*** *Difference in event-related spectral perturbation between left and right STN during linguistic inhibition (left) and motor inhibition (right).*

***Supplementary Table 1.*** *Patient characteristics. UPDRS = Universal Parkinson's Disease Rating scale, LEDD = Levodopa equivalent dose, MoCA = Montreal Cognitive Assessment*

| | | | | | | | | Left STN DBS | | | | Right STN DBS | | | |
|---|---|---|---|---|---|---|---|---|---|---|---|---|---|---|---|
| **#** | **Age** | **Disease Duration (Y.)** | **Hoehn-Yahr** | **UPDRS On** | **UPDRS Off** | **LEDD** | **MoCA** | **Hz** | **Voltage** | **Pulse Width (uS)** | **Polarity** | **Hz** | **Voltage** | **Pulse Width (uS)** | **Polarity** |
| 1 | 69 | 9 | 2 | 14 | 27 | 220 | 24 | 125/125 | 2.2/0.5 | 60/60 | Int-mono | 125/125 | 2.2/0.5 | 60/60 | Int-mono |
| 2 | 58 | 5 | 3 | 34 | 65 | 688 | 23 | 155 | 0.7 | 60 | Mono | 155 | 2.2 | 60 | Mono |
| 3 | 73 | 11 | 4 | 38 | 64 | 500 | 22 | 120 | 1.35 | 60 | Mono | 120 | 4 | 60 | Mono |
| 4 | 53 | 5 | 2 | 18 | 50 | 250 | 29 | 130 | 2.8 | 60 | Bipolar | 130 | 2.35 | 60 | Bipolar |
| 5 | 59 | 9 | 2 | 22 | 65 | 375 | 26 | 130 | 2.05 | 100 | Mono | 130 | 2.25 | 80 | Mono |
| 6 | 68 | 4 | 2 | 35 | 59 | 188 | 27 | 135 | 2.4 | 60 | Mono | 135 | 1.8 | 60 | Mono |
| 7 | 49 | 13 | 3 | 28 | 57 | 500 | 29 | 130 | 3.35 | 80 | Mono | 130 | 3.35 | 80 | Mono |
| 8 | 71 | 9 | 2 | 33 | 68 | 500 | 29 | 130 | 1.35 | 60 | Bipolar | 130 | 3.1 | 60 | Bipolar |
| 9 | 73 | 9 | 1 | 14 | 36 | 0 | 26 | 165 | 6 | 60 | Mono | 165 | 0.8 | 60 | Mono |
| 10 | 61 | 3 | 2 | 27 | 56 | 500 | 26 | 145 | 1.3 | 60 | Mono | 145 | 1.3 | 70 | Mono |
| 11 | 61 | 12 | 2 | 26 | 48 | 0 | 26 | 160 | 2.55 | 70 | Mono | 160 | 2.6 | 60 | Mono |
| 12 | 71 | 14 | 4 | 45 | 66 | 1375 | 25 | 95 | 3.25 | 60 | Mono | 95 | 3.3 | 60 | Mono |
| 13 | 68 | 12 | 3 | 14 | 46 | 1875 | 26 | 145 | 1.6 | 70 | Mono | 145 | 1.65 | 70 | Mono |
| 14 | 66 | 2 | 3 | 41 | 72 | 0 | 22 | 130 | 1.8 | 60 | Bipolar | 130 | 2.6 | 60 | Bipolar |
| 15 | 73 | 8 | 3 | 44 | 59 | 1250 | 23 | 150 | 2.4 | 60 | Mono | 150 | 2.3 | 70 | Mono |
| 16 | 65 | 6 | 2 | 36 | 46 | 813 | 24 | 140 | 2.7 | 60 | Mono | 140 | 3.5 | 60 | Mono |
| 17 | 72 | 8 | 2 | 38 | 64 | 125 | 27 | 175 | 4.3 | 20 | Mono | 175 | 3.3 | 60 | Mono |
| 18 | 80 | 6 | 2 | 31 | 53 | 250 | 23 | 185 | 2.3 | 100 | Mono | 185 | 2.1 | 60 | Mono |
| 19 | 67 | 7 | 2 | 33 | 51 | 500 | 27 | 125 | 2.4 | 60 | Bipolar | 125/125 | 2.9/1 | 60/90 | Int-bi |
| 20 | 57 | 3 | 2 | 8 | 29 | 1000 | 25 | 130 | 0.4 | 60 | Mono | 130 | 3.6 | 70 | Mono |

**Supplementary Table 2.** DBS system lead model and MNI reconstruction information

| Subject | Lead model | Pair | mni_x | mni_y | mni_z |
|---|---|---|---|---|---|
| 1 | 3387 | R6/R8 | 12.6735 | -9.33574 | -2.7353915 |
| 2 | 3387 | R6/R8 | 9.950595 | -9.010795 | -5.33988 |
| 3 | B33015 | R10-12/R16 | 11.124375 | -10.61337667 | -4.626376667 |
| 4 | B33015 | R9/R13-15 | 10.6603333 | -11.92681667 | -8.18498 |
| 5 | B33015 | R9/R13-15 | 11.8443667 | -12.37396667 | -8.487613333 |
| 6 | 3387 | R6/R8 | 8.849155 | -7.067125 | -5.84699 |
| 7 | B33015 | R10-12/R16 | 10.341725 | -10.55326 | -3.5050745 |
| 8 | B33015 | R10-12/R16 | 9.15199167 | -10.31146833 | -5.645576667 |
| 9 | B33015 | R10-12/R16 | 10.0737533 | -9.686628333 | -3.6698895 |
| 10 | B33015 | R9/R16 | 10.750305 | -11.72087 | -6.896145 |
| 11 | B33015 | R9/R13-15 | 11.46545 | -11.55742333 | -7.347663333 |
| 12 | B33015 | R10-12/R16 | 8.387575 | -10.13274333 | -2.8401515 |
| 13 | B33015 | R10-12/R16 | 11.6581167 | -11.84665 | -4.595535 |
| 14 | B33015 | R9/R13-15 | 12.24205 | -12.24283333 | -6.68792 |
| 15 | B33015 | R10-12/R16 | 10.756855 | -12.92226667 | -2.534104 |
| 16 | 3387 | R5/R7 | 13.08835 | -13.10375 | -7.673835 |
| 17 | 3387 | R5/R7 | 15.09515 | -14.5076 | -6.003115 |
